# PKMζ-KIBRA interactions, molecular turnover, and memory

**DOI:** 10.64898/2026.01.05.697418

**Authors:** Changchi Hsieh, David A. Cano, Panayiotis Tsokas, James E. Cottrell, André Antonio Fenton, Harel Shouval, Todd Charlton Sacktor

**Affiliations:** Department of Physiology and Pharmacology, The Robert F. Furchgott Center for Neural and Behavioral Science, State University of New York Downstate Health Sciences University; Brooklyn, New York 11203, USA; College of Medicine, State University of New York Downstate Health Sciences University; Brooklyn, New York 11203, USA; Department of Anesthesiology, State University of New York Downstate Health Sciences University; Brooklyn, New York 11203, USA; Department of Pathology, State University of New York Downstate Health Sciences University; Brooklyn, New York 11203, USA; Center for Neural Science, New York University; New York, New York 10003, USA; Neuroscience Institute at NYU Langone Medical Center; New York, New York 10016, USA; Department of Neurobiology and Anatomy, University of Texas Medical at Houston, Houston, TX 77030, USA; Department of Neurology, State University of New York Downstate Health Sciences University; Brooklyn, New York 11203, USA

**Keywords:** PKMzeta, PKM-zeta, long-term potentiation (LTP), WWC1

## Abstract

How can the molecules that strengthen synaptic connections maintain memory in the face of molecular turnover? Our previous work showed that persistent interaction between the postsynaptic scaffolding protein, KIBRA, and the autonomously active PKC isoform, PKMζ, is crucial for maintaining synaptic long-term potentiation (LTP) and memory for at least a month. This duration is longer than the lifespans of individual KIBRA and PKMζ molecules. Biophysical modeling of the interaction suggests oligomers of KIBRA-PKMζ dimers, but not individual dimers or monomers, can overcome molecular turnover by continually incorporating newly synthesized KIBRA and PKMζ, replacing those that have degraded. Here we used AlphaFold 3 to predict the structures of KIBRA-PKMζ heterodimers and heterohexamers and to examine the sites of action of two structurally distinct inhibitors of KIBRA-PKMζ interaction that disrupt established late-LTP and long-term memory. The structures predict that the peptide K-ZAP blocks formation of heterodimers, whereas the small molecule ζ-stat prevents PKMζ of one heterodimer from binding a second KIBRA and PKMζ, essential for forming larger oligomeric structures. We show that ζ-stat, like K-ZAP, disrupts 1-month-old spatial memory. Thus, continual formation of KIBRA-PKMζ oligomers can be a core molecular mechanism driving the persistence of long-term memory in the face of molecular turnover.

## Introduction

How can memories be maintained when all molecular components of synapses continually turn over [1]? Autonomously active protein kinases can maintain synaptic potentiation, but they turn over within hours to days [2]. Crick and Lisman independently proposed a mechanism to prolong the action of these kinases for the lifespan of a memory— although the molecules themselves are not long-lived, the interactions between them can persist while newly synthesized proteins continually replace those that degrade [1, 2].

Two kinases that potentiate synaptic transmission become autonomously active [3]. One is Ca^2+^/calmodulin-dependent kinase II (CaMKII) that becomes Ca^2+^/calmodulin-independent through autophosphorylation and is crucial for the initial stages of inducing LTP and memory [4]. The other is the continually active atypical PKC, PKMζ, that is critical for sustaining the mechanistically distinct, enduring maintenance of wild-type LTP and memory [5]. Once translated [6], the steady-state increase in PKMζ persists for hours to maintain late-LTP in hippocampal slices and for at least a month during spatial memory in specific synaptodendritic regions of the hippocampal neurons that were active during learning [7, 8].

To maintain LTP and memory PKMζ must continuously interact with a synaptic tag, the postsynaptic scaffolding protein KIBRA (aka WWC1), which is genetically linked to human memory performance and Alzheimer’s disease [9, 10]. After initial synthesis, KIBRA and PKMζ form complexes that persist during maintenance and continually target the kinase’s action to active synapses [10] (Fig. 1a). Importantly, two structurally distinct inhibitors disrupt KIBRA-PKMζ interaction and reverse established late-LTP and long-term spatial memory [10]. One, the peptide inhibitor K-ZAP mimics and occludes the action of a sequence in KIBRA’s C-terminus where PKMζ binds [11]. The other, the small molecule ζ-stat blocks the “handle” motif of PKMζ’s catalytic domain where KIBRA binds [10, 12].

**Fig 1.**
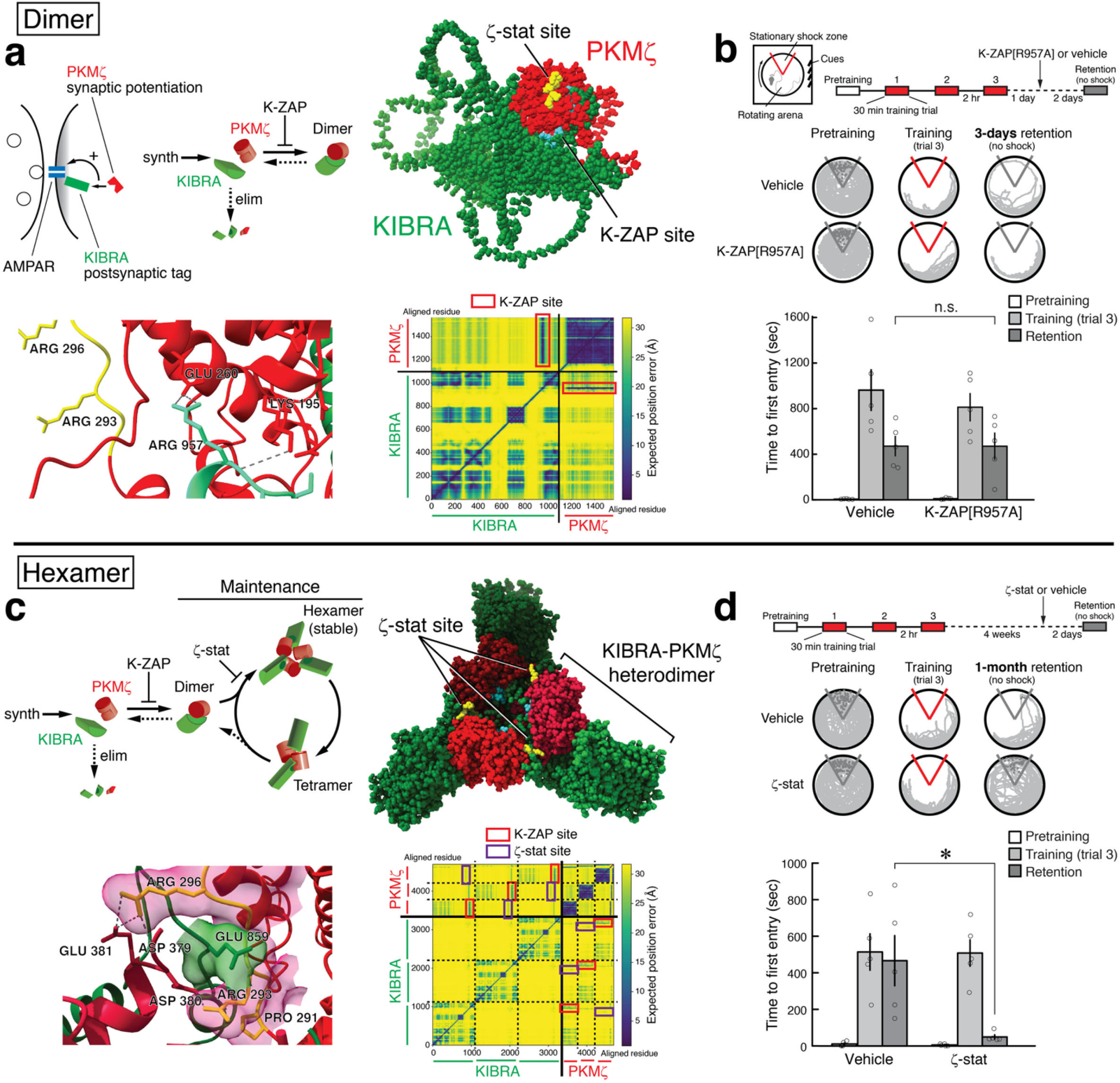
KIBRA-PKMζ interactions predicted to form dimers and hexamers maintain long-term spatial memory. **a** Above left, schematic showing functional elements of KIBRA-PKMζ synaptic potentiation. Postsynaptic KIBRA acts as a synaptic tag that targets PKMζ, which potentiates AMPAR responses. Above middle, summary of kinetic model of formation of KIBRA-PKMζ heterodimers from new synthesis (synth) of monomers. High levels of KIBRA-tags (green) and PKMζ (red) cannot be maintained because dimers dissociate, and the elimination (elim) of monomers is rapid. Above right, heterodimer structure predicted by AlphaFold 3. KIBRA’s K-ZAP sequence (blue) interacts with PKMζ, whereas the PKMζ’s handle does not (ζ-stat binding site, i.e., 7 amino acids from P291 to F297, shown in yellow). Below left, the 3 hydrogen bonds between K-ZAP arginine-957 and PKMζ are shown; there are 20 hydrogen bonds between the total K-ZAP sequence and PKMζ. The arginines in the ζ-stat-binding motif of PKMζ do not interact with KIBRA (yellow, amino-acid numbering based on PKMζ sequence [6]). Below right, predicted aligned error plot of the KIBRA-PKMζ heterodimer. K-ZAP sequence interaction with PKMζ are shown in red boxes. **b** Whereas K-ZAP disrupts 1-day- and 1-month-old spatial memory [10] (shown in kinetic model, **a** above middle), bilateral hippocampal injections of inactive, mutated K-ZAP peptide (myristoyl-FVRNSLE**A**RSVRMKRPS, 5 nmol in 0.5 μl vehicle per side) does not disrupt 1-day-old memory. Above left, schematic of active place avoidance training apparatus shows a slowly rotating arena containing a nonrotating shock zone sector (delineated in red). Visual cues located on the walls of the room are needed to avoid the shock zone. Above right, protocol for active place avoidance. Middle, representative paths during 10 min of pretraining, at end of training trial 3, and 1-day memory retention. Below, mean ± SEM. ANOVA with repeated measurements reveals a single significant effect of training (pretraining, training, and retention; *F*_2,16_ = 33.81, *P* < 0.00001, *η*^2^_p_ = 0.81), and no treatment effect (vehicle and mutated K-ZAP[R957A]) nor their interaction. Bonferroni-corrected comparisons confirm that the memory retention after K-ZAP[R957A] injection does not significantly differ from vehicle control (n.s., *P* = 1; as initial experiments with K-ZAP[R957A] showed normal memory, for comparison with K-ZAP[R957A], vehicle controls were pooled with 4 randomly selected vehicle controls from K-ZAP experiments [10], n’s = 5). **c** Above left, kinetic model showing self-perpetuating formation of stable KIBRA-PKMζ hexamers in LTP/memory maintenance. Above right, predicted hexamer structure. KIBRA’s K-ZAP sequence interacts with PKMζ forming pairs, and PKMζ’s handle links the pair to a second KIBRA (dark green) and PKMζ (dark red). Below left, PKMζ’s ζ-stat binding site interacts with a second KIBRA (interaction shown with molecular surfaces) and forms 4 hydrogen bonds with a second PKMζ. Below right, predicted aligned error plot of KIBRA-PKMζ hexamers. K-ZAP sequence interaction sites with PKMζ are shown in red boxes, ζ-stat sites in purple. **d** Bilateral hippocampal injections of ζ-stat (5 nmol in 0.5 μl vehicle per side) disrupt 4-week-old spatial memory. Above, protocol for active place avoidance. Middle, representative paths during 10 min of pretraining, at end of training trial 3, and 4-week memory retention. Below, mean ± SEM. ANOVA with repeated measurements finds significant effects of training (pretraining, training, and retention; *F*_2,62_ = 25.93, *P* = 0.00001, *η*^2^_p_ = 0.76) and interaction between effects of training and treatment (vehicle and ζ-stat) (training X treatment: *F*_2,16_ = 5.79, *P* = 0.01, *η*^2^_p_ = 0.42). The 1-month memory retention was abolished by ζ-stat injected 2 days prior to the test, compared with vehicle (*, significant Tukey *post-hoc* tests, *P* = 0.008, n’s = 5).

It is crucial that PKMζ and KIBRA interact; however, biophysical modeling suggests that simple PKMζ-KIBRA heterodimers cannot permanently store information at synapses [13]. PKMζ monomers rapidly degrade and become highly stable upon binding KIBRA [12-14]. Nevertheless, heterodimers formed after LTP induction or learning will eventually dissociate into rapidly degrading monomers. Consequently, the information they once encoded is lost.

Our biophysical model predicts that hexameric or larger oligomers composed of KIBRA-PKMζ pairs are better suited to store information over long time periods because they can survive molecular turnover [13]. Specifically, individual degraded molecules of the larger complexes can be replaced because the remaining oligomer can serve as a template for binding newly synthesized KIBRA and PKMζ. This view predicts that the inhibitors K-ZAP or ζ-stat that disrupt KIBRA-PKMζ interactions would prevent the continual replenishment of the oligomers and reverse synapses from a stable potentiated to stable unpotentiated state. This outcome would permanently disrupt late-LTP and long-term memory, a result that has been observed [10].

## Results

For heterodimers to form larger complexes, one molecule of each species should bind to more than one molecule of the other. To investigate if KIBRA and PKMζ have this property, we used AlphaFold 3 to predict both their dimeric and hexameric forms.

### The K-ZAP sequence of KIBRA that binds PKMζ forms KIBRA-PKMζ dimers

AlphaFold 3 predicts that heterodimers are formed by interaction between the K-ZAP sequence of KIBRA (FVRNSLERRSVRMKRPS-966), and the surface of PKMζ (Fig. 1a). However, the PKMζ-handle, where ζ-stat binds, does not appear to interact with KIBRA in the dimeric complex.

The peptide K-ZAP disrupts 1-day- and 1-month-old memory [10]. To further test whether K-ZAP interaction is critical for maintaining memory, we trained mice on an active place avoidance memory task and 1 day later injected bilaterally into hippocampus a mutated form of K-ZAP, in which the critical KIBRA arginine-957 is changed to alanine to decrease the peptide’s interaction with PKMζ [14] (Fig. 1b). In the predicted heterodimers, arginine-957 has 3 hydrogen bonds with PKMζ, and the mutation to alanine (K-ZAP[R957A]) has only 1. If K-ZAP prevents memory maintenance by interfering with PKMζ-KIBRA dimerization, then the mutated version should have no effect. As predicted, hippocampal injections of the mutated peptide K-ZAP[R957A] did not affect long-term memory retention.

### The PKMζ-handle with the ζ-stat binding site interacts with a second KIBRA and PKMζ in KIBRA-PKMζ hexamers

In hexamers, KIBRA’s K-ZAP sequence preserves KIBRA-PKMζ pairing, and the PKMζ-handle binds to a second KIBRA as well as a second PKMζ, linking the pairs (Fig. 1c). Two amino acids in the PKMζ handle, proline-291 and phenylalanine-297, are critical for both strong binding of PKMζ to KIBRA and the inhibitory action of ζ-stat [10]. These amino acids flank two arginines predicted to interact with a disordered region of KIBRA and the surface of another PKMζ. If these flanking amino acids are changed to the analogous amino acids of the other atypical PKCι/λ, which binds only weakly to KIBRA, the mutated PKMζ[PKCι/λ-P291Q;F297S] also binds weakly to KIBRA [10]. On changing the flanking amino acids in the structural model, the predicted number of hydrogen bonds linking the PKMζs decreases from 4 to 0 [10]. Consequently, AlphaFold 3 predicts that the ζ-stat-binding site is key to forming and maintaining the hexamers. The site is precisely where the KIBRA-PKMζ pairs interact with each other. This contrasts with dimers in which the ζ-stat-binding site does not participate.

The estimated lifespans of individual PKMζ and KIBRA molecules is a few days [10]. We found that injecting ζ-stat to inhibit the predicted KIBRA-PKMζ oligomer-interaction site disrupts a 4-week-old spatial memory (Fig. 1d). Thus, 1-month long-term memory depends on KIBRA-PKMζ oligomers for its maintenance.

## Discussion

Biophysical modeling of KIBRA-PKMζ interaction predicted that hetero-oligomers could maintain high levels of the KIBRA-tag and PKMζ at active synapses to sustain potentiation despite protein turnover [13]. Therefore, if ζ-stat specifically prevents oligomer formation as AlphaFold predicts (Fig. 1c), then, like K-ZAP that blocks dimer formation [10], ζ-stat should disrupt memories that are maintained longer than the lifespans of individual PKMζ and KIBRA molecules. This was observed (Fig. 1d). A critical feature of our kinetic model is that the formation of hexamers from dimers should include a cooperative step [13]. The putative binding of a PKMζ to two KIBRAs and a second PKMζ might provide the nonlinearity necessary to produce hexamers. Our simple model does not exclude the possibility that other molecules are important components of KIBRA-PKMζ complexes, such as PICK1 that can interact with both KIBRA and PKMζ [5, 15]. Characterizing the core mechanisms for the self-perpetuation of KIBRA-PKMζ complexes with their associated proteins might elucidate the fundamental molecular properties of a synaptic “mnemosome” that stores information in the brain and is disrupted in disorders of memory.

## Abbreviations

AMPAR: α-amino-3-hydroxy-5-methyl-4-isoxazolepropionic acid receptor;
CaMKII: Ca^2+^/calmodulin-dependent kinase
II; KIBRA: KIdney BRAin protein;
K-ZAP: KIBRA-PKMζ antagonist peptide;
LTP: long-term potentiation;
PICK1: protein interacting with C-kinase 1;
PKCι/λ: protein kinase C iota/lambda;
PKCζ: protein kinase C zeta;
PKMζ: protein kinase Mzeta;
WWC1: WW and C2 Domain Containing protein 1

## Acknowledgments

The authors declare no financial interests.

## Author contributions

Conceptualization: TCS, AAF

Methodology: CH, DAC

Investigation: CH, DAC

Visualization: CH, DAC

Funding acquisition: TCS, AAF, JEC

Project administration: TCS, AAF

Supervision: TCS, AAF

Writing – original draft: TCS, AAF

Writing – review & editing: TCS, AAF, PT, JEC, CH, DAC

## Declarations

### Ethics approval and consent to participate

Not applicable.

This study was performed in strict accordance with the recommendations in the Guide for the Care and Use of Laboratory Animals of the National Institutes of Health. All animals were handled according to approved Institutional Animal Care and Use Committee (IACUC) protocols [no. 11-10274, 15-10467 of the State University of New York (SUNY) Downstate Health Sciences University; animal welfare assurance number: D16-00167].

### Consent for publication

Not applicable.

## Materials and Methods

### Protein modeling

FASTA protein sequences for KIBRA (Q5SXA9) and PKMζ (Q02956-2) from *Mus musculus*, were taken from Uniprot and analyzed by AlphaFold 3 to generate protein complexes *in silico*. All models contained one ATP and two Mg^2+^ ions per PKMζ molecule, as well as activating post-translational modifications for PKMζ at P-threonine-227 and P-threonine-377 (corresponding to P-threonine 410 and P-threonine 560 in PKCζ). The highest confidence AlphaFold 3 output files were visualized in UCSF ChimeraX (v1.9). Hydrogen bonds were calculated by ChimeraX with distance tolerance set to 0.4 Å and angle tolerance set to 20°.

### Active Place Avoidance Conditioning

All experiments were performed blindly. Active place avoidance and intrahippocampal injections were performed as previously described [10]. Briefly, active place avoidance was conducted with a commercial computer-controlled system (Bio-Signal Group, Acton, MA). The mouse was placed on a 40-cm diameter circular arena rotating at 1 rpm. The specialized software, Tracker (Bio-Signal Group, Acton, MA), was used to detect the animal’s position 30 times per second by video tracking from an overhead camera. The time to first enter the shock zone estimates ability to avoid shock and was taken as an index of between-session long-term place avoidance memory. The training schedule was as follows: after a 30-min pretraining session, the animals received three 30-min training trials, with an intertrial interval of 2 hours. Long-term memory retention was tested either 3 days or 30 days later without shock. The drugs were administered 2 days before the retention test. Pre-established exclusion criterion was if cannulae were found to be incorrectly targeted. No mice were excluded.

### Statistics

Multi-factor comparisons were performed using mixed-design ANOVA with repeated measures or Bonferroni-corrected *t*-tests, as appropriate. The degrees of freedom for the *F* values of the ANOVAs are reported as subscripts. *Post-hoc* multiple comparisons were performed by Tukey tests as appropriate. Statistical significance was accepted at *P* < 0.05. Effect sizes for multi-factor ANOVAs are reported as *η*^2^_p_.

